# KAP is the neuronal organelle adaptor for Kinesin-2 KIF3AB and KIF3AC

**DOI:** 10.1101/2022.07.30.502069

**Authors:** Alex Garbouchian, Andrew Montgomery, Susan P. Gilbert, Marvin Bentley

**Author notes:** Correspondence to: Marvin Bentley, Rensselaer Polytechnic Institute, 110 8th Street, Troy, NY, 12180.

## Abstract

Kinesin-driven organelle transport is crucial for neuron development and maintenance, yet the mechanisms by which kinesins specifically bind their organelle cargoes remain undefined. In contrast to other transport kinesins, the neuronal function and specific organelle adaptors of heterodimeric Kinesin-2 family members KIF3AB and KIF3AC remain unknown. We developed a novel microscopy-based assay to define protein–protein interactions in intact neurons. The experiments found that KIF3AB and KIF3AC both bind KAP. These interactions are mediated by the distal C-terminal tail regions and not the coiled-coil domain. We used live-cell imaging in cultured hippocampal neurons to define the localization and trafficking parameters of KIF3AB and KIF3AC organelle populations. We discovered that KIF3AB/KAP and KIF3AC/KAP bind the same organelle populations and defined their transport parameters in axons and dendrites. The results also show that ∼12% of KIF3 organelles contain the RNA binding protein, adenomatous polyposis coli. These data point towards a model in which KIF3AB and KIF3AC use KAP as their neuronal organelle adaptor and that these kinesins mediate transport of a range of organelles.

## INTRODUCTION

The Kinesin-2 family consists of heterodimeric and homodimeric processive motors that mediate microtubule plus-end directed transport (Yamazaki et al., 1995; Yang and Goldstein, 1998; Kondo et al., 1994; Muresan et al., 1998; Scholey, 2013; Guzik-Lendrum et al., 2015; Gilbert et al., 2018). Mammals express three heterodimeric Kinesin-2 proteins: KIF3A, KIF3B, and KIF3C, that assemble into two dimers, KIF3AB or KIF3AC. KIF3AB is the primary anterograde motor in intraflagellar transport (IFT) (Funabashi et al., 2018; Milic et al., 2017; Engelke et al., 2019; Prevo et al., 2017; Lewis et al., 2018; Scholey, 2013), and binds IFT particles through the adaptor kinesin-associated protein 3B (KAP) (Wedaman et al., 1996; Yamazaki et al., 1996; Brunnbauer et al., 2010; Carpenter et al., 2015; Scholey, 2013; Funabashi et al., 2018).

Beyond the well-defined role of KIF3AB/KAP in IFT, little is known about its other transport functions. KIF3AB/KAP is proposed to mediate transport of spectrin/fodrin (Takeda et al., 2000) and RNA particles containing adenomatous polyposis coli (APC) (Ruane et al., 2016; Jimbo et al., 2002). The latter function was recently supported by elegant in vitro experiments showing that KIF3AB/KAP and APC assemble into a transport-capable complex, although neuronal evidence of this interaction is still outstanding (Baumann et al., 2020). Even less is known about KIF3AC. KIF3C is highly expressed in the central nervous system (Navone et al., 2001; Muresan et al., 1998; Sardella et al., 1998) and cultured hippocampal neurons (Silverman et al., 2010), but its neuronal function is unclear. To date, the only proposed KIF3AC cargo is fragile-X mental retardation protein (FMRP) (Davidovic et al., 2007), and KIF3CC homodimers may mediate microtubule dynamics in response to axon injury (Gumy et al., 2013). Recent studies made some progress towards characterizing KIF3AB and KIF3AC in neurons. Constitutively active motor domain heterodimers of KIF3AB and KIF3AC can move on both axonal and dendritic microtubules (Huang and Banker, 2012). However, motor domain activity does not necessarily reflect transport preferences of kinesins that are associated with their vesicle cargoes (Nabb et al., 2020). Beyond these initial studies, little is known about the neuronal function of KIF3AB and KIF3AC, raising the following questions: do KIF3AB and KIF3AC participate in neuronal organelle transport, what is the adaptor for organelle–cargo linkage, and what other proteins or motors associate with the KIF3 organelles in hippocampal neurons?

To address these questions, we applied two primary strategies in cultured hippocampal neurons. First, we developed a fluorescence-based assay to probe for protein–protein interactions in intact neurons. We systematically characterized the binding interactions of KIF3AB and KIF3AC with KAP and found that both KIF3AB and KIF3AC bind KAP. Moreover, the KAP binding site for both heterodimers is localized in their C-terminal disordered domains. Second, we expressed fluorescent kinesin tail domain in cultured hippocampal neurons to analyze the localization and trafficking parameters of organelle-bound kinesins (Montgomery et al., 2022; Yang et al., 2019). With this approach, we found that KIF3AB, KIF3AC, and KAP bind the same neuronal organelles. These organelles undergo sporadic long-range transport. A fraction of these organelles colocalize with the RNA-binding protein adenomatous polyposis coli, indicating that KIF3/KAP likely interacts with additional neuronal cargoes. Results show that KIF3AB/KAP and KIF3AC/KAP participate in neuronal transport outside of IFT and that KAP is an organelle adaptor for KIF3AB and KIF3AC.

## RESULTS

### A novel microscopy-based protein–protein interaction assay reveals that KAP binds KIF3AB and KIF3AC

KIF3AB interaction with KAP is well established in IFT (Verhey et al., 2011; Yamazaki et al., 1996; Brunnbauer et al., 2010; Gilbert et al., 2018). In contrast, the function of KIF3AB and KIF3AC outside IFT is not well understood, especially in neurons. One key unknown is whether KIF3AB and KAP assemble in neurons outside of IFT. A second unknown is whether KIF3AC binds KAP. To determine KIF3–KAP interactions, we developed a fluorescence-based protein– protein binding assay in intact neurons (Fig. 1A). Biochemical binding assays have long been the standard in the cell biological toolkit for determining protein–protein interactions. While powerful, we sought to determine interactions in their natural milieu, the cytoplasm of cultured hippocampal neurons, a highly specialized cell type (Bentley and Banker, 2016; Yogev and Shen, 2017; Arendt et al., 2016, 2019).

**Figure 1.**
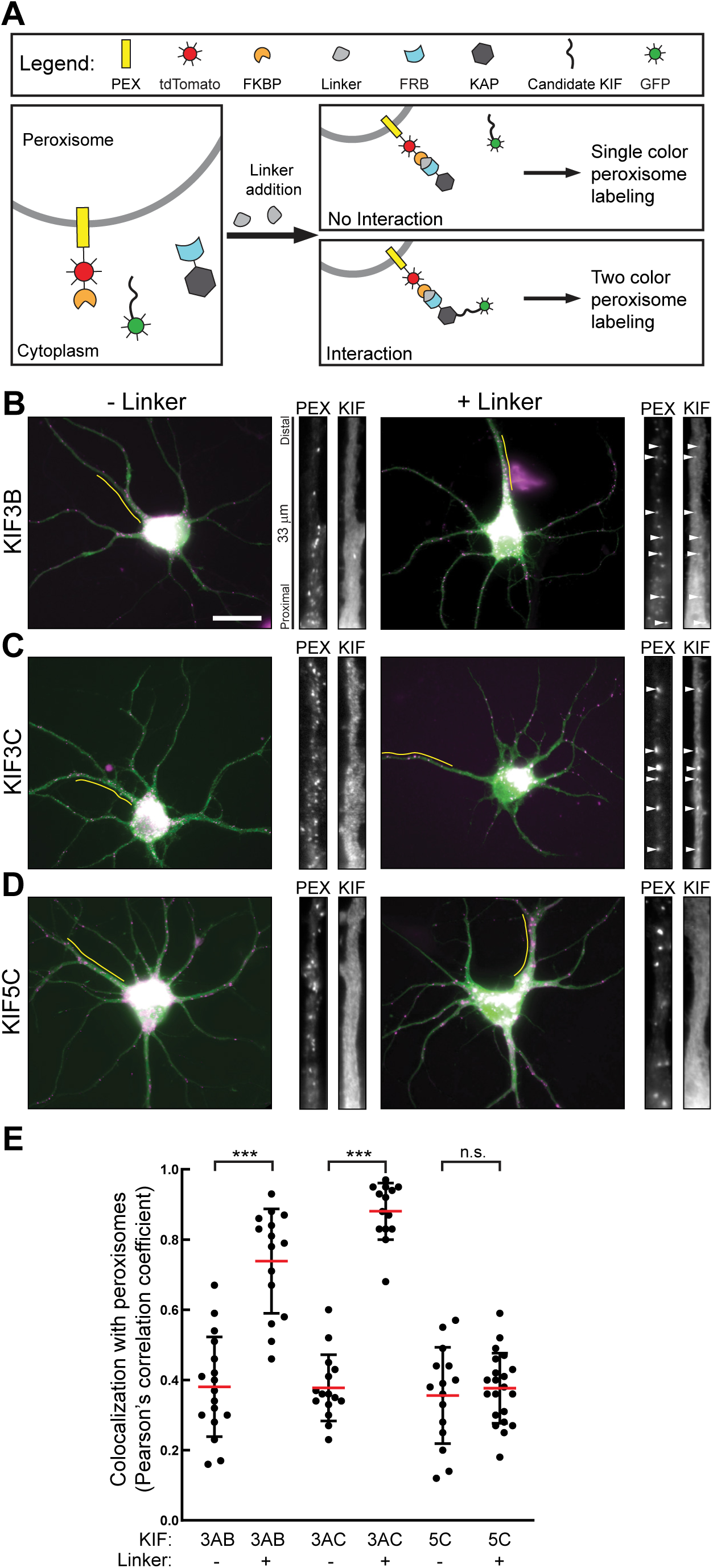
KIF3AB and KIF3AC each bind KAP. (A) A schematic illustrating the protein–protein interaction assay. Four proteins are expressed together: PEX-tdTM-FKBP, FRB-KAP, unlabeled KIF3A tail, and a GFP-tagged kinesin tail. Binding between KAP and the kinesin is evaluated by linker-induced colocalization of the GFP-tagged KIF3 tail with peroxisomes. (B-D) Representative images of 7 DIV hippocampal neurons expressing the indicated constructs. High magnification images of dendrites were taken from the regions indicated by the yellow line. Arrowheads highlight examples of colocalization. Scale bar: 20 µm. (E) Quantification of colocalization between peroxisomes and kinesins. Cell sample sizes: KIF3AB(-): 15; KIF3AB(+): 15; KIF3AC(-): 15; KIF3AC(+): 14; KIF5C(-): 15; KIF5C(+): 21. ***p < 0.001.

The assay was designed to use inducible colocalization as the readout for binding between two proteins, in this case KAP and the candidate KIF3s. It involves expression of proteins tagged with the FK506 binding protein (FKBP) and FKBP12-rapamycin binding (FRB) domains, which dimerize in the presence of a rapamycin analog linker molecule (Belshaw et al., 1996; Robinson et al., 2010; Jenkins et al., 2012; Bentley and Banker, 2015; Bentley et al., 2015; Banaszynski et al., 2005; Kapitein et al., 2010) (Fig. 1A). Candidate GFP-tagged kinesin tails and FRB-KAP were coexpressed with a construct consisting of an FKBP domain, a tdTomato fluorophore, and the peroxisomal protein Pex3 (PEX-tdTM-FKBP). Peroxisomes are convenient organelles because they are a distinct and largely stationary organelle population that is not thought to interact with KIF3s. Linker-specific colocalization of GFP-KIF3 and peroxisomes indicates specific interaction between FRB-KAP and the GFP-tagged candidate kinesin tail.

We first applied the assay to test for interactions between FRB-KAP and GFP-KIF3B^391-747^ tail (Fig. 1B), as there is a strong prediction for binding of these proteins. Both proteins were coexpressed with untagged KIF3A tail and PEX-tdTM-FKBP. High magnification images show that PEX-tdTM-FKBP was targeted to peroxisomes in dendrites, as anticipated (Kapitein et al., 2010). GFP-KIF3B was mostly soluble in the absence of linker, and there was no colocalization with peroxisomes. In cells treated with 100 nM linker for 2 h, the distribution of peroxisomes was unchanged. In contrast, linker treatment resulted in GFP-KIF3B colocalization with peroxisomes. Not all GFP-KIF3B was targeted to peroxisomes; a soluble component remained in the cytoplasm. This is presumably because peroxisomal targeting is limited by the availability of FRB-KAP and PEX-tdTM-FKBP. However, any linker-specific colocalization is extraordinarily strong evidence that KIF3B and KAP specifically interact in neurons.

We next asked if KIF3C tail (GFP-KIF3C^411-796^) binds KAP in neurons (Fig. 1C). Without linker, GFP-KIF3C was mostly soluble and did not colocalize with peroxisomes. Treatment with linker targeted GFP-KIF3C to peroxisomes, indicating that KAP bound KIF3C. This is the first experimental evidence to show specific KAP–KIF3C interaction. This finding also suggests that KAP is a neuronal KIF3 adaptor outside of IFT, because KIF3C cannot rescue IFT transport in cilia when KIF3B is not expressed (Engelke et al., 2019).

Finally, we tested GFP-KIF5C tail (GFP-KIF5C^378-955^) to confirm the specificity of the assay (Fig. 1D). KIF5C is a member of the Kinesin-1 family that does not interact with KAP (Schnapp, 2003; Hackney et al., 1991; Woźniak and Allan, 2006; Verhey et al., 2001). As expected, linker treatment did not target GFP-KIF5C to peroxisomes, which confirms the specificity of the assay.

Because the readout of these experiments is colocalization of the candidate kinesin and peroxisomes, the Pearson’s correlation coefficient is a convenient quantification method (Fig. 1E). Without linker, correlation coefficients were consistently between 0.3 and 0.4, typical values for cellular structures and a largely soluble protein that does not specifically colocalize (Frank et al., 2020). Linker addition and the resulting colocalization caused high coefficients for KIF3B (0.75) and KIF3C (0.88). The correlation coefficient for KIF5C was not changed by linker treatment. Together, these measurements show that KAP binds not just KIF3AB but also KIF3AC, and that this assay is highly specific.

### KIF3 tails dimerize without the N-terminal coiled-coil dimerization domain

The protein–protein binding assay allowed us to further investigate the KIF3AB/KAP and KIF3AC/KAP interactions. First, we asked if KIF3 tails dimerize (Fig. 2A-D). These constructs lack the N-terminal motor, neck linker, and first coiled-coil domains that are sufficient for dimerization and generate constitutively active processive motors (Huang and Banker, 2012; Guzik-Lendrum et al., 2015; Phillips et al., 2016; Guzik-Lendrum et al., 2017; Gilbert et al., 2018; Andreasson et al., 2015). GFP-KIF3B^391-747^ (Fig. 2C) or GFP-KIF3C^411-796^ (Fig. 2D) were coexpressed with FRB-KIF3A and FKBP-tdTM-PEX. Linker treatment caused both GFP-tagged tails to colocalize with peroxisomes, which was accompanied by an increase in the Pearson’s correlation coefficient (Fig. 2E). These data show that KIF3AB and KIF3AC tails form dimers independent of the N-terminal motor and coiled-coil dimerization domains.

**Figure 2.**
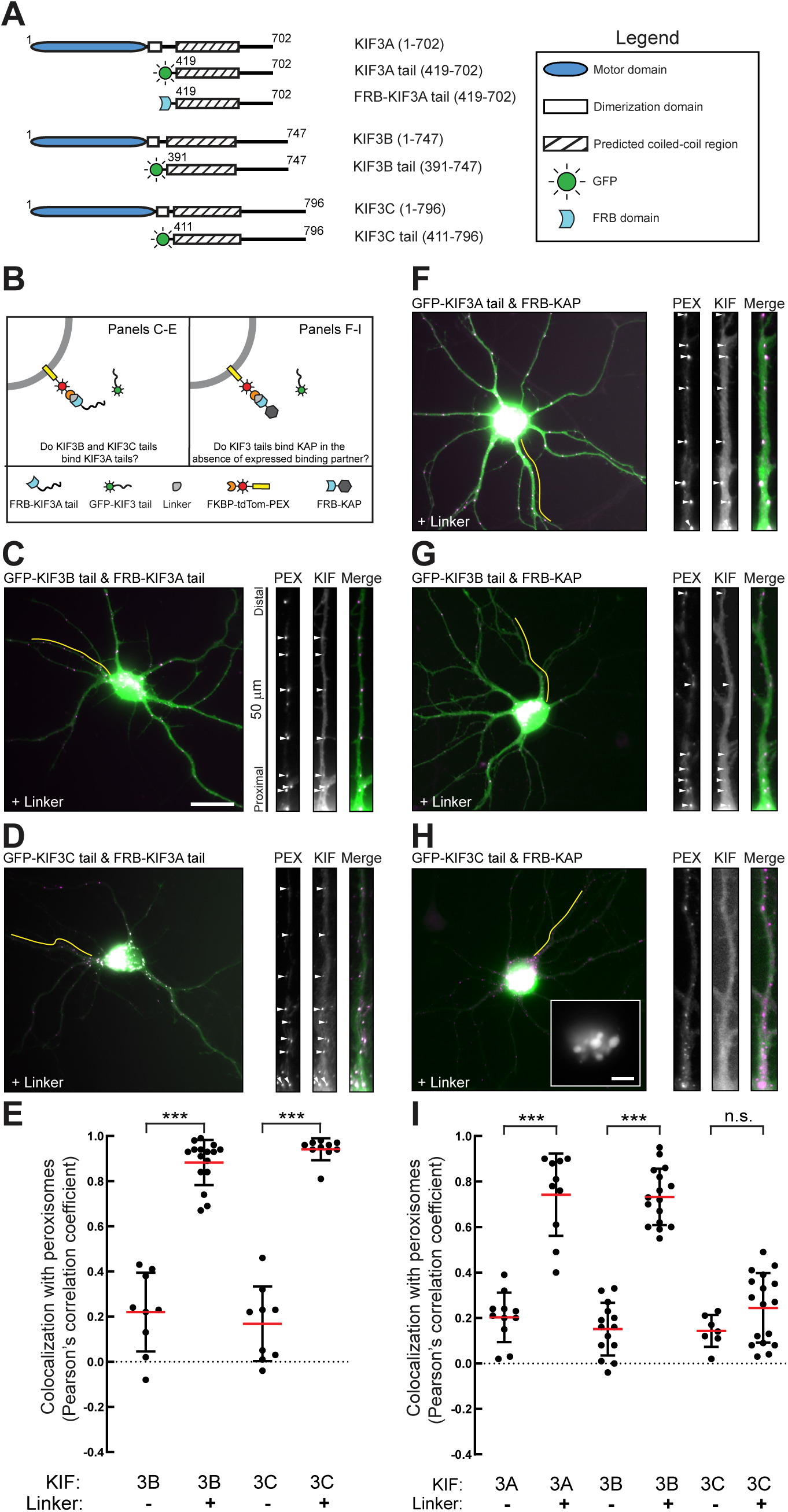
KIF3AB and KIF3AC dimerization does not require the canonical N-terminal dimerization domain. (A) A schematic showing the construct design. (B) A schematic illustrating the specific application of the protein–protein binding assay. (C&D, F-H) Representative images of 9 DIV neurons expressing the indicated proteins and treated with linker. High magnification images of dendrites were taken from the regions indicated by the yellow line. Arrowheads highlight examples of colocalization. The inset in H shows GFP-KIF3C aggregates in the soma. Low magnification scale bar: 20 µm. High magnification inset scale bar: 5 µm. (E&I) Quantification of colocalization between peroxisomes and kinesins. Cell sample sizes panel E: KIF3B(-): 10; KIF3B(+): 16; KIF3C(-): 9; KIF3C(+): 11. Cell sample size panel I: KIF3A(-): 11; KIF3A(+): 10; KIF3B(-): 14; KIF3B(+): 16; KIF3C(-): 7; KIF3C(+): 17. ***p < 0.001.

The fact that KIF3AB and KIF3AC tails form heterodimers may indicate that such dimerization is a prerequisite for binding KAP. Because kinesins are typically in a stable dimeric state, exogenously expressed tails are unlikely to dimerize with endogenous kinesins. Therefore, expressing one KIF3 tail without its dimerization partner will likely result in monomers that could not bind KAP if dimerization is a prerequisite for that interaction. To test this hypothesis, we expressed GFP-KIF3A, GFP-KIF3B, or GFP-KIF3C tail together with FRB-KAP and PEX-tdTM-FKBP. GFP-KIF3A (Fig. 2F) and GFP-KIF3B (Fig. 2G) both colocalized with peroxisomes after linker treatment, indicating that they can bind KAP as monomers. Quantification by Pearson correlation coefficient corroborated these results (Fig. 2I). In contrast, GFP-KIF3C (Fig. 2H) clustered in large protein aggregates in the soma instead of the mostly soluble distribution it exhibits when coexpressed with KIF3A (Fig. 2D). Improperly folded proteins commonly form aggregates, and the fact that aggregates only appeared when GFP-KIF3C tail was expressed without KIF3A tail suggests that monomeric KIF3C does not fold properly. Therefore, KIF3CC homodimers are unlikely to form during normal development, although there may be exceptions to this in response to axon injury (Gumy et al., 2013).

### Disordered KIF3 C-terminal domains bind KAP

KIF3 tails are divided into two structural domains: a predicted coiled-coil domain and a C-terminal disordered domain (Delorenzi and Speed, 2002; Gilbert et al., 2018; Yamazaki et al., 1995; Wedaman et al., 1996). Early electron microscopy studies proposed that KAP binds the C-terminal domain of KIF3AB (Yamazaki et al., 1996; Wedaman et al., 1996). In contrast, recent studies in *Drosophila* found that KAP binds the coiled-coil domain of heterodimeric KIF3 (Doodhi et al., 2009; Ahmed et al., 2020). Because KIF3AC–KAP interactions have not been reported prior to the present study, there are no proposed models for KIF3AC–KAP binding interactions.

To identify the KAP binding domains of KIF3A, KIF3B, and KIF3C, we generated fluorescently tagged expression constructs of the coiled-coil (KIF3^CC^, Fig. 3A) or disordered (KIF3^DD^, Fig. 4A) domains and determined if each bound KAP. Because coexpression may be required for stable expression of KIF3, we first expressed each coiled-coil with its binding partner. We coexpressed GFP-KIF3A^CC^ and Halo-KIF3B^CC^ with FRB-KAP and PEX-tdTM-FKBP (Fig. 3C). In a separate experiment, we coexpressed GFP-KIF3A^CC^ and Halo-KIF3C^CC^ with FRB-KAP and PEX-tdTM-FKBP (Fig. 3D). Halotags were visualized with JF646. Colocalization of each coiled-coil with peroxisomes was then quantified (Fig. 3E). None of the coiled-coils were targeted to peroxisomes by linker addition, indicating that none of the coiled-coils interacted with KAP.

**Figure 3.**
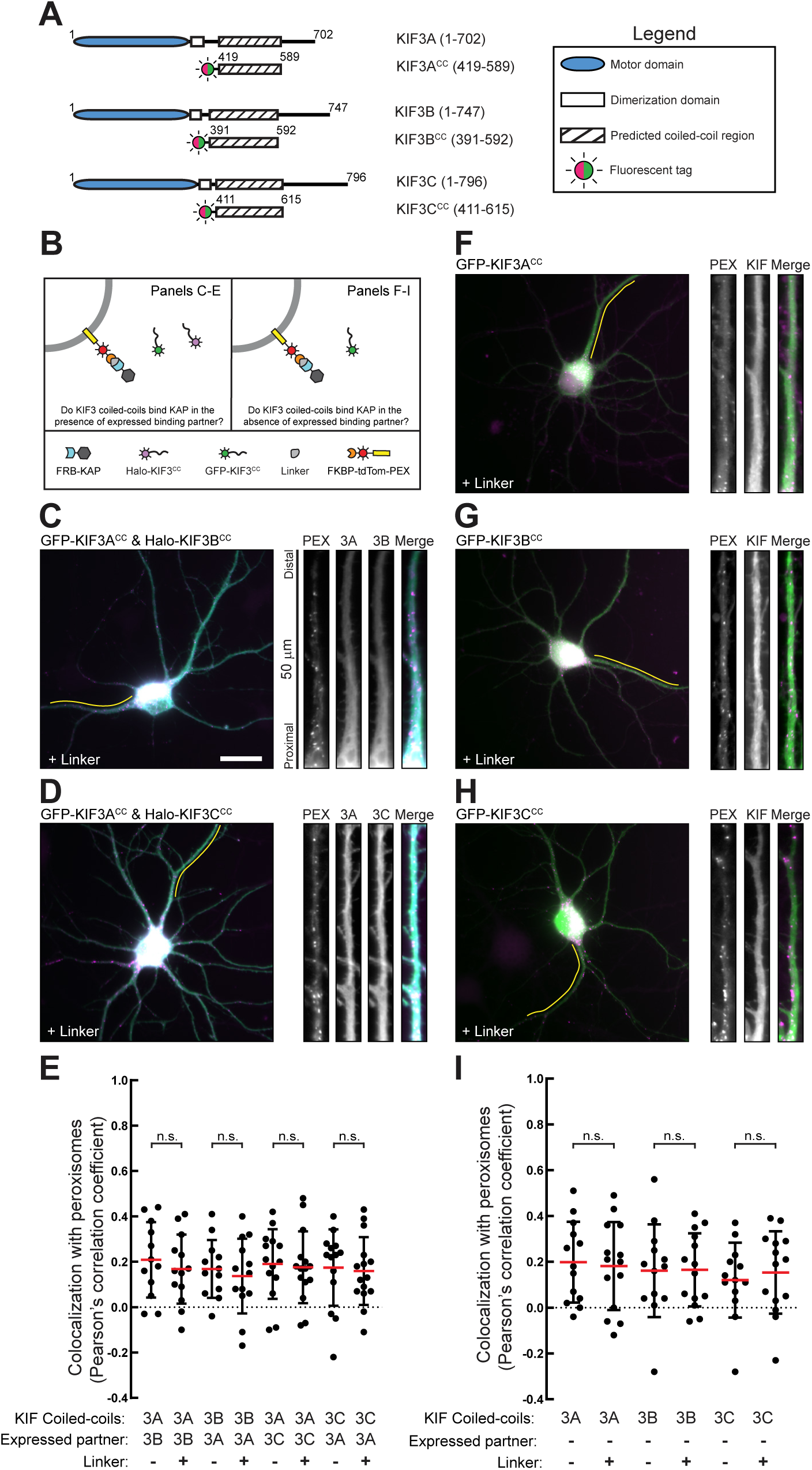
KAP binding to KIF3 is not mediated by coiled-coil domains. (A) A schematic showing the construct design. (B) A schematic illustrating the specific application of the protein–protein binding assay. (C&D, F-H) Representative images of 7 DIV neurons expressing the indicated proteins and treated with linker. High magnification images of dendrites were taken from the regions indicated by the yellow line. Scale bar: 20 µm. (E&I) Quantification of colocalization between peroxisomes and kinesins. Cell sample sizes panel E: KIF3AB^CC^(-): 12; KIF3AB^CC^(+): 15; KIF3AC^CC^(-): 14; KIF3AC^CC^(+): 15. Cell sample sizes panel I: KIF3A^CC^(-): 13; KIF3A^CC^(+): 14; KIF3B^CC^(-): 13; KIF3B^CC^(+): 14; KIF3C^CC^(-): 13; KIF3C^CC^(+): 14. n.s. not significant.

**Figure 4.**
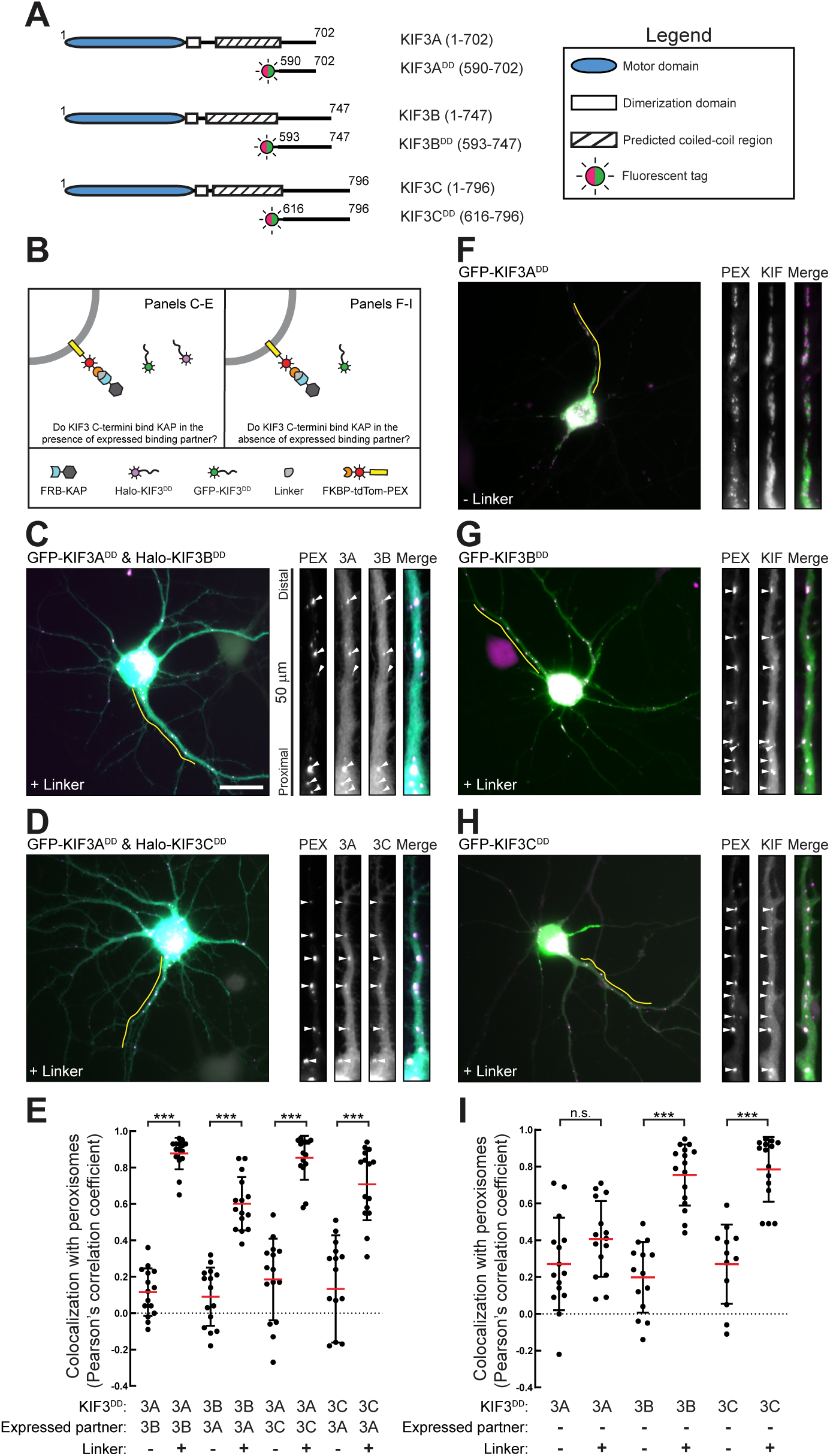
KAP binds the KIF3 C-terminal disordered domain. (A) A schematic showing the construct design. (B) A schematic illustrating the specific application of the protein–protein binding assay. (C&D, F-H) Representative images of 7 DIV neurons expressing the indicated proteins and treated with linker. High magnification images of dendrites were taken from the regions indicated by the yellow line. Arrowheads highlight examples of colocalization. Scale bar: 20 µm. (E&I) Quantification of colocalization between peroxisomes and kinesins. Cell sample sizes panel E: KIF3AB^DD^(-): 15; KIF3AB^DD^(+): 15; KIF3AC^DD^(-): 15; KIF3AC^DD^(+): 15. Cell sample sizes panel I: KIF3A^DD^(-): 15; KIF3A^DD^(+): 15; KIF3B^DD^(-): 14; KIF3B^DD^(+): 16; KIF3C^DD^(-): 12; KIF3C^DD^(+): 15. ***p < 0.001.

We next asked if KIF3 coiled-coils behaved differently when expressed without their binding partner. Each coiled-coil (GFP-KIF3A^CC^, GFP-KIF3B^CC^, or GFP-KIF3C^CC^) was coexpressed with FRB-KAP and PEX-tdTM-FKBP (Fig. 3F-H). All coiled-coil constructs expressed without forming aggregates. Linker treatment did not result in colocalization of coiled-coil domains with peroxisomes. Corresponding Pearson’s correlation coefficients confirmed that linker treatment did not cause colocalization with peroxisomes (Fig. 3I). Together, these results are strong evidence that mammalian KIF3 coiled-coils do not interact with KAP.

The finding that the coiled-coil domains of mammalian KIF3s do not interact with KAP predicts that the disordered C-terminal domain mediates interaction with KAP. We used the same systematic strategy to determine if the disordered KIF3 domains bind KAP (Fig. 4A&B). We coexpressed GFP-KIF3A^DD^ and Halo-KIF3B^DD^ with FRB-KAP and PEX-tdTM-FKBP (Fig. 4C). Next, we coexpressed GFP-KIF3A^DD^ and Halo-KIF3C^DD^ with FRB-KAP and PEX-tdTM-FKBP (Fig. 4D). Colocalization of each KIF3 disordered domain with peroxisomes was quantified by Pearson’s correlation coefficient (Fig. 4E). All three KIF3 disordered domains were targeted to peroxisomes in linker-treated neurons, showing that these domains alone are sufficient for binding KAP. A limitation of this experiment is that it cannot determine if individual KIF3^DD^ bind to KAP, or if KIF3AB^DD^ and KIF3AC^DD^ bind KAP as dimers. To address this, we coexpressed each disordered domain with FRB-KAP and PEX-tdTM-FKBP (Fig. 4F-H). Colocalization of KIF3 disordered domains with peroxisomes was quantified as before (Fig. 4I). We first noticed that GFP-KIF3A^DD^ did not express properly. Instead of the expected soluble distribution, GFP-KIF3A^DD^ formed aggregates in the somatodendritic region. Aggregate formation was independent of linker incubation and aggregates did not form when KIF3A^DD^ was coexpressed with either KIF3B^DD^ or KIF3C^DD^ (Fig. 4C-E). Therefore, the C-terminal disordered domains may dimerize independent of the coiled-coil domain. This is surprising, as coiled-coil domains are typically the primary sites of kinesin dimerization. In contrast, GFP-KIF3B^DD^ and GFP-KIF3C^DD^ were distributed through the cytoplasm without linker, and without any aggregate formation. Both colocalized with peroxisomes after linker treatment. This shows that KIF3B^DD^ and KIF3C^DD^ bind KAP, and that this binding interaction does not depend on dimerization.

### KIF3AB, KIF3AC, and KAP organelles exhibit similar cellular distributions and neuronal transport parameters

Because KIF3B and KIFC both bind KAP, they may interact with some of the same neuronal organelle populations. We recently developed techniques for visualizing organelle-bound kinesins in live cells (Montgomery et al., 2022; Yang et al., 2019; Frank et al., 2020). Improved organelle labeling was achieved by expressing the GFP-tagged kinesin tail domains, which mediate organelle binding. Using this approach we generated initial recordings of organelle-bound KIF3B and KIF3C in neurons (Yang et al., 2019). However, the signal-to-noise in these experiments was low and may have missed any subpopulation of organelles that bound fewer kinesins, as these would be dim and challenging to image. It also prevented quantitative analysis of KIF3B and KIF3C organelle transport parameters.

To quantitatively assess the localization and transport parameters of KIF3AB and KIF3AC organelles, we sought to improve the technique. To facilitate recording at early expression times and improve visualization of dim organelles, we used the halotag system (Los et al., 2008; Encell, 2012). The halotag enzyme itself is not fluorescent and becomes visible only after covalent binding to a fluorescent ligand. Bright organic dyes have substantially more favorable fluorescence characteristics than fluorescent proteins (Grimm et al., 2015; Xia et al., 2013) and enhance detection of organelle-bound proteins (Montgomery et al., 2022; Frank et al., 2020; Yang et al., 2019). To use this approach, we generated halo-tagged KIF3B and KIF3C tail constructs (Fig. 5A).

**Figure 5.**
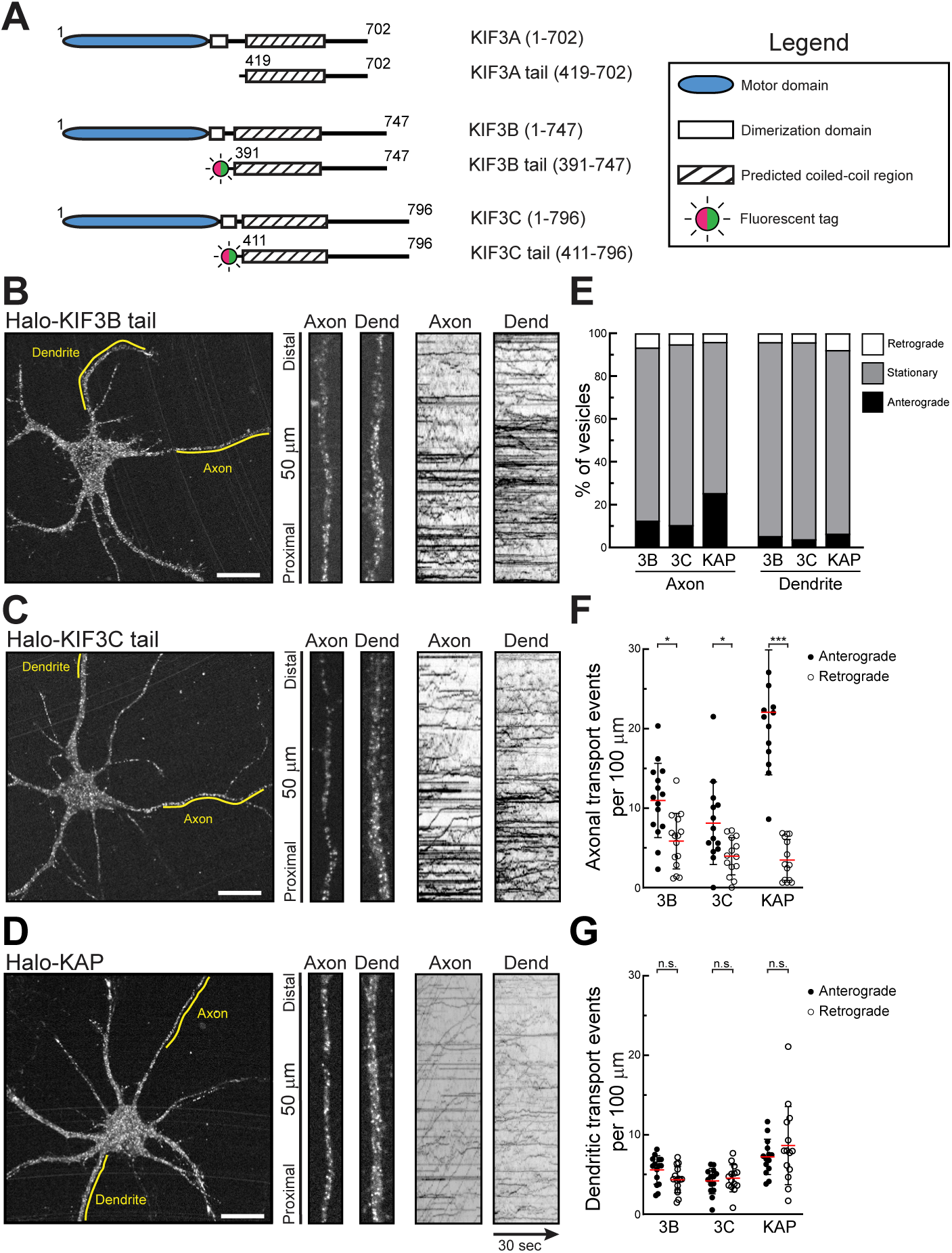
Characterization of KIF3B, KIF3C, and KAP organelles in cultured hippocampal neurons. (A) A schematic showing the construct design. (B-D) Representative images of 7-8 DIV hippocampal neurons expressing the indicated proteins. GFP-KIF3B and GFP-KIF3C were coexpressed with untagged KIF3A tail. Halotag was visualized with JF549. Yellow lines on each cell indicate sections of the axon and dendrite from which the high magnification views and kymographs were generated. (E-G) Quantification of organelle motility and transport events. Sample sizes: KIF3B: 15 cells, 1,027 axonal and 4,741 dendritic organelles; KIF3C: 15 cells, 1,090 axonal and 4,747 dendritic organelles; KAP: 14 cells, 1,001 axonal and 4,470 dendritic organelles. *p<0.05; ***p<0.001.

We expressed Halo-KIF3B (Fig. 5B) or Halo-KIF3C (Fig. 5C) tail in cultured hippocampal neurons and visualized halotag with JF549 (Grimm et al., 2015). Because these kinesin tails form KIF3AB or KIF3AC dimers, we coexpressed non-fluorescent KIF3A tail to ensure the dimerization partner was present at similar expression levels. This approach resulted in detectable organelles as early as 3 h after transfection.

Halo-KIF3B organelles were distributed throughout the neurons. High magnification images from the axon and a dendrite show that organelles were consistently round and less than 1 µm in diameter. Corresponding kymographs show the movement of Halo-KIF3B organelles. On a kymograph, lines with a positive slope indicate anterograde movement and lines with a negative slope indicate retrograde movement. This convention is followed in all kymographs. Long-range transport events of Halo-KIF3B organelles were infrequent, and most did not extend beyond 5 µm.

Halo-KIF3C organelles exhibited a localization and morphology comparable to that of Halo-KIF3B (Fig. 5C). Organelles were in both axon and dendrites, with infrequent long-range movements. One difference to KIF3B was that Halo-KIF3C tail expression consistently produced less cytoplasmic background than Halo-KIF3B. This higher signal-to-noise eased detection but did not result in a total increase in the number of labeled organelles.

Because both KIF3B and KIF3C bind KAP, these results suggest that KAP participates in the transport of neuronal KIF3AB and KIF3AC organelles. KAP is a globular protein with few known binding partners (Yamazaki et al., 1996; Carpenter et al., 2015), and little is known about its neuronal localization. We expressed Halo-KAP and visualized organelles with JF549 (Fig. 5D). Halo-KAP labeled organelles in axons and dendrites. Most organelles had a round morphology, with a diameter <1 µm, comparable to that of KIF3B and KIF3C organelles. In the axon, KAP organelles underwent long-range transport with a strong anterograde bias. Halo-KAP expression resulted in very little cytoplasmic background fluorescence and consistent labeling.

Overall, KIF3B, KIF3C, and KAP organelles exhibited multiple commonalities. They were homogenously distributed in axons and dendrites, exhibited comparable morphology, and underwent sporadic transport. Notably, despite differences in cytoplasmic background and the associated differences in single-to-noise, all three constructs labeled roughly the same number of organelles (see legend to Fig. 5).

Achieving consistency in organelle labeling in combination with live-cell imaging allowed us to quantitatively assess the transport characteristics of each population. Each organelle in representative dendrite and axon segments was scored for its motility (Fig. 5E). In the axon, most KIF3B, KIF3C, and KAP organelles were stationary. Retrograde movement was rare, exhibited by fewer than 7 % of organelles. All three populations exhibited some amount of anterograde axonal movement. KAP exhibited the most active anterograde transport (25 %), followed by KIF3B (12 %) and KIF3C (10 %). In dendrites the three populations behaved nearly identically. Movements were infrequent and without any directional bias. To evaluate directionality biases in transport, we determined the relative frequency of anterograde and retrograde movements in axons and dendrites by kymograph analysis (Fig. 5F&G). In axons, all three populations displayed an anterograde bias, although the bias was most pronounced for KAP. For all organelles, there were fewer overall transport events in dendrites. In dendrites, transport exhibited no directional bias, which is consistent with the fact that dendritic microtubules are less densely packed and have a mixed orientation (Baas et al., 1988; Tas et al., 2017).

Transport parameters differ between populations and act as a quantitative “signature” for specific organelles (Frank et al., 2020; Yang et al., 2019; Burack et al., 2000; Niwa et al., 2008; Koppers and Farías, 2021; Bodakuntla et al., 2020; De Pace et al., 2020; Lo et al., 2011; Farfel-Becker et al., 2019; Bommel et al., 2019). We systematically quantified the transport parameters (i.e., velocity and run length) of the three organelle populations (Fig. 6). Histograms show run length and velocity measurements of KIF3B, KIF3C, and KAP moving anterograde or retrograde in axons or dendrites (Fig. 6A-D). KIF3B and KIF3C organelles behaved nearly identically. One difference between KIF3B and KIF3C organelles was in retrograde axonal movements, although this was likely due to variability caused by the relatively small number of total organelles undergoing retrograde movement in axons.

**Figure 6.**
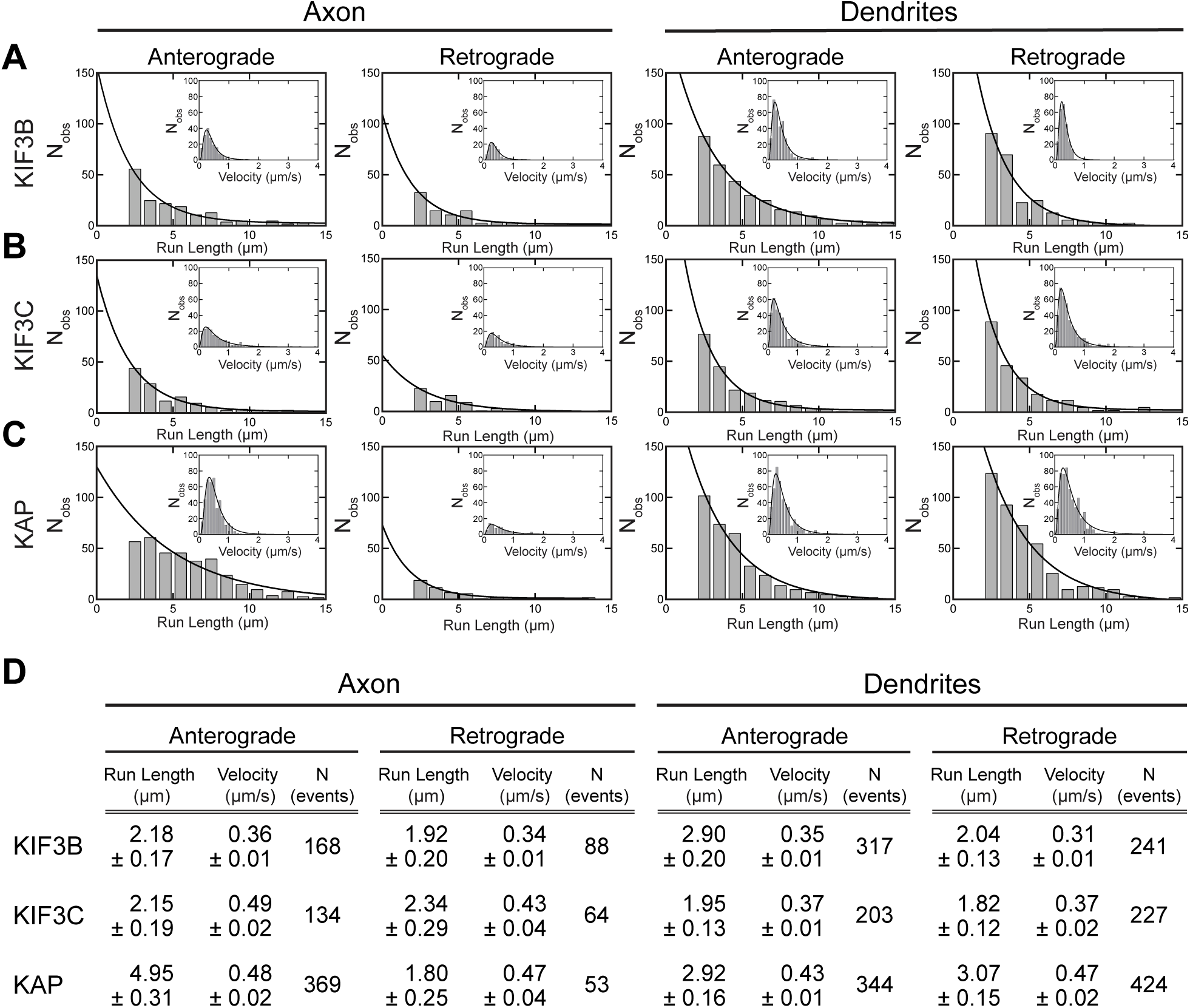
KIF3B, KIF3C, and KAP organelles exhibit comparable transport parameters. (A-C) Histograms showing the distribution of organelle run lengths and velocities with SEM. (D) Compiled transport parameters. Mean run length was generated by fitting to a single-exponential decay function. Cell sample sizes: KIF3B: 15; KIF3C: 14; KAP: 14.

Mean organelle velocities, ranging from 0.31 to 0.49 µm/s, were slower than organelle populations that are moved by other kinesins (Yang et al., 2019; Nabb and Bentley, 2022; Niwa et al., 2008; Takeda et al., 2000), but were comparable to the previously reported velocities for KIF3AB and KIF3AC in single-molecule motility experiments (Guzik-Lendrum et al., 2015; Gilbert et al., 2018; Bensel et al., 2017; Woll et al., 2018). KAP organelles consistently exhibited comparable velocities to KIF3B and KIF3C. However, the mean anterograde run length of KAP organelles was 4.95 µm compared to 2.18 µm for KIF3B and 2.15 µm for KIF3C in axons.

There are at least three plausible explanations for this result. First, KAP may bind to some organelles without KIF3AB or KIFAC. This could be either because KAP acts as an adaptor for other kinesins or serves some other function on these organelles. There is little evidence supporting this, as there are no other proposed functions of KAP. A second possible explanation is that the kinesin tail strategy could cause a dominant negative effect and reduce transport. However, the comparable velocities between KAP and KIF3AB/KIF3AC suggests that the motor complexes are not substantially impaired in their ability to move organelles. Furthermore, previous experiments with kinesin tail constructs did not find any substantial change in the trafficking behavior of cargo-labeled vesicles (Jenkins et al., 2012; Yang et al., 2019; Montgomery et al., 2022; Frank et al., 2020; Bentley and Banker, 2015). A third interpretation is that KAP binding to organelles is more stable than KIF3AB or KIF3AC binding to KAP. If KAP mediates KIF3 binding to organelles, KAP detachment would also result in the loss of bound kinesins. In contrast, if kinesin–KAP interactions were severed, KAP could remain on moving organelles, even if the kinesin signal was lost. This model is consistent with the observation that KAP expression typically resulted in higher signal-to-noise organelle labeling with less cytoplasmic background. Furthermore, KAP can form a complex with endogenous kinesins, whereas expressed KIF3AB and KIF3AC rely on endogenous adaptors, likely KAP, to bind to organelles. It is clear from the dim fluorescent signal that a small number of kinesins and KAP bind to any given organelle at once. If kinesins cycle on and off organelles at a higher rate, it is plausible that fluorescent kinesins detach during movement. This is consistent with the larger number of moving KAP organelles we observed.

Overall, these results show that KIF3B, KIF3C, and KAP organelles displayed comparable transport parameters, size and morphology, and cellular distribution. The similarities in their distribution and their transport parameters suggest the hypothesis that some fraction of the organelles labeled by KIF3B, KIF3C, and KAP are identical.

### KIF3AB, KIF3AC, and KAP bind the same neuronal organelle populations

To determine the amount of KIF3AB, KIF3AC, and KAP colocalization on the same organelles, we performed a series of two-color experiments (Fig. 7). We expressed GFP-KIF3B tail with Halo-KAP (Fig. 7A), GFP-KIF3C tail with Halo-KAP (Fig. 7B), or GFP-KIF3B tail with Halo-KIF3C tail (Fig. 7C), each coexpressed with untagged KIF3A. Halotags were visualized with JF549 prior to fixation and imaging. High magnification images from representative dendrites show that there was overwhelming overlap in all three conditions. We quantified colocalization by measuring Pearson’s correlation coefficients in dendritic regions with organelles (Fig. 7D). For cellular imaging data, no correlation typically results in Pearson’s correlation coefficients lower than 0.4. All three conditions resulted in coefficients of 0.8 or higher, indicating extraordinarily high colocalization. Together, these data indicate KIF3B, KIF3C, and KAP bind the same organelle populations.

**Figure 7.**
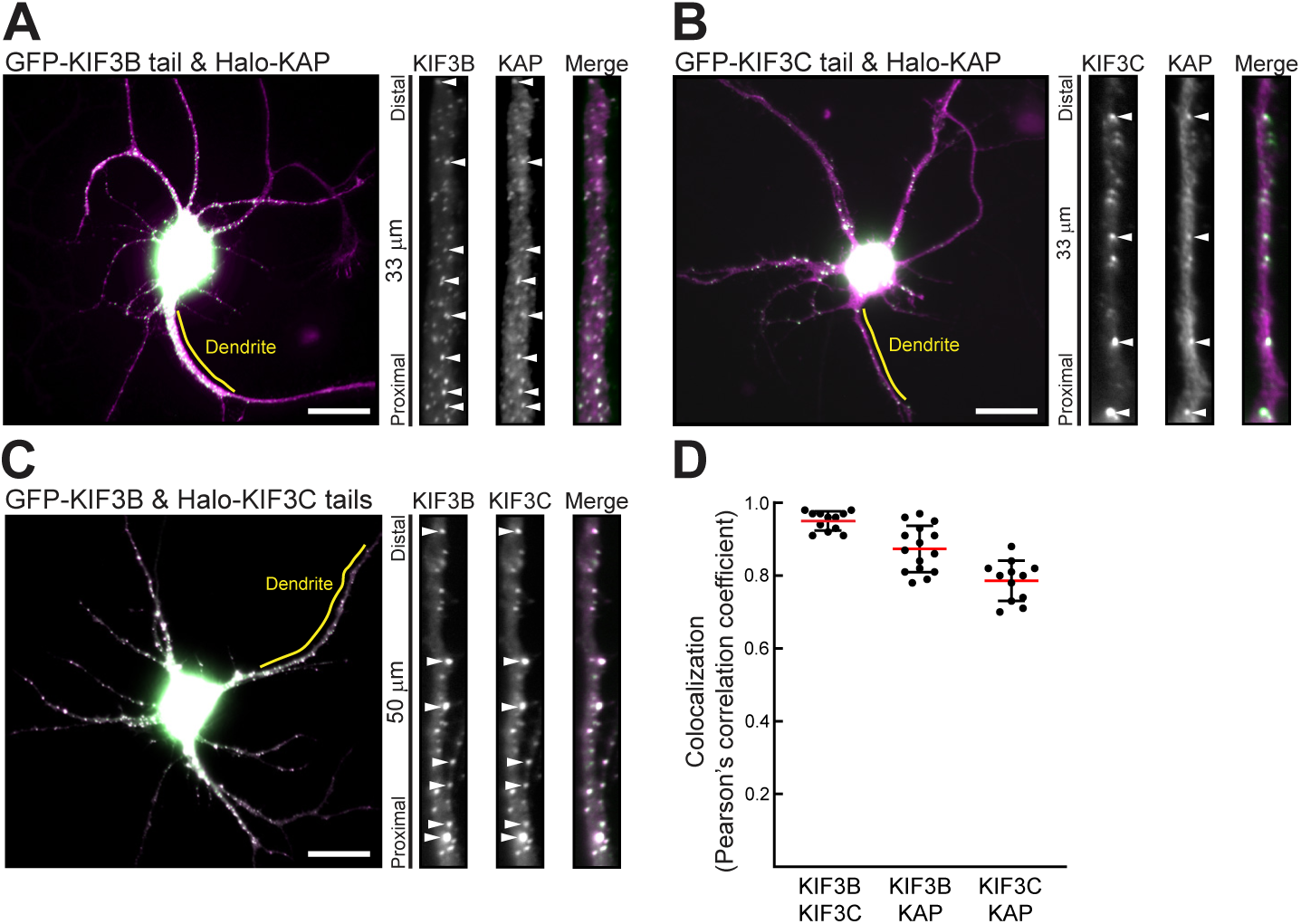
KIF3B, KIF3C, and KAP bind the same neuronal organelle populations. (A-C) Representative images of 7-9 DIV hippocampal neurons expressing the indicated constructs. KIF3B and KIF3C were coexpressed with untagged KIF3A tail. Yellow lines indicate regions from which high magnification views were generated. Arrowheads point to examples of colocalization. Scale bar: 20 µm. (D) Quantification of colocalization of the indicated proteins. Cell sample sizes: KIF3B/KIF3C: 12; KIF3B/KAP: 15; KIF3C/KAP: 12.

### Adenomatous polyposis coli is a neuronal KIF3 cargo

Beyond IFT, little is known about neuronal KIF3 cargoes. The strongest evidence of a non-IFT cargo is adenomatous polyposis coli (APC), although this interaction is not well characterized in neurons (Alsabban et al., 2020; Baumann et al., 2020; Jimbo et al., 2002). There is also neuronal immunofluorescence evidence that KIF3C binds to organelles containing fragile X mental retardation protein (FMRP) (Davidovic et al., 2007). Consistent colocalization of fluorescently tagged proteins, especially when colocalization remains stable for longer periods in live cells, is some of the strongest evidence that two proteins bind the same organelles. Because KIF3B and KIF3C bind the same organelles, we chose KIF3C tail, which labeled organelles more consistently, and determined colocalization with fluorescently tagged APC and FMRP. We used the KIF5 cargo neuron-glia cell adhesion molecule (NgCAM; L1 in mammals) (Yang et al., 2019; Nabb and Bentley, 2022; Jareb and Banker, 1998; Sampo et al., 2003; Lasiecka et al., 2010; Hoo et al., 2016) as a baseline for incidental colocalization. We expressed Halo-KIF3C (visualized with JF549) and unlabeled KIF3A tails with APC-mEmerald, GFP-FMRP, or NgCAM-GFP (Fig. 8A-C). High magnification images show that APC colocalized with KIF3C (Fig. 8A). In contrast, KIF3C did not colocalize with FMRP or NgCAM (Fig. 8B&C). We quantified the percentage of KIF3C organelles colocalized with each cargo as well as the percentage of cargo that colocalized with KIF3C (Fig. 8D). About 12 % of KIF3C structures colocalized with APC in axons and dendrites. KIF3C rarely colocalized with FMRP and NgCAM. Therefore, APC is a likely KIF3 cargo. These results also suggest that, despite previous data, FMRP is likely not a KIF3AC cargo in hippocampal neurons. The quantification also indicates that KIF3 binds to cargoes other than APC that are not yet identified.

**Figure 8.**
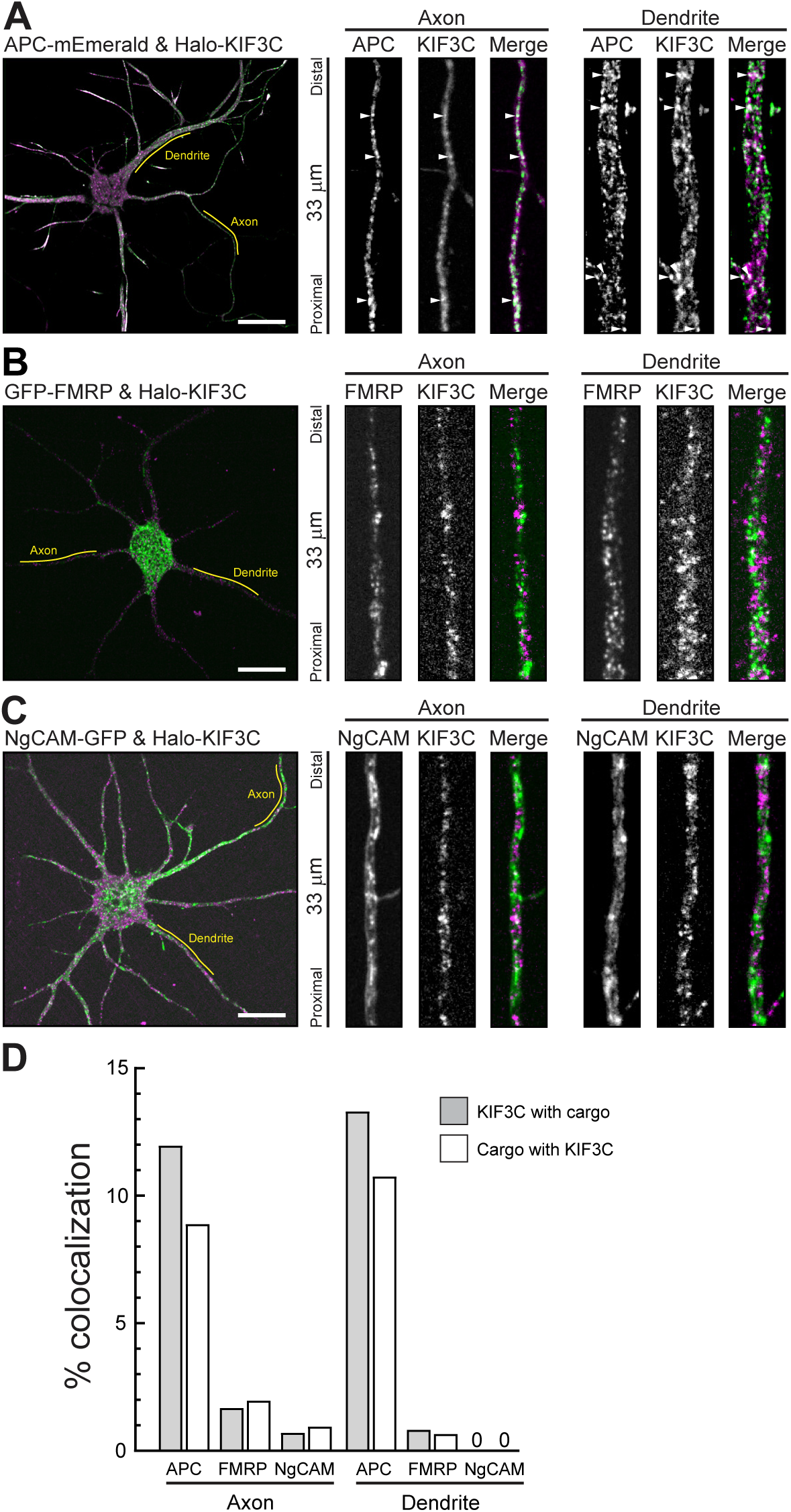
KIF3/KAP colocalizes with APC and not FMRP or NgCAM. (A-C) Representative images of 6-9 DIV hippocampal neurons expressing the indicated constructs and untagged KIF3A tail. Halotags were visualized with JF549. Yellow lines indicate regions from which high magnification views were generated. Arrowheads highlight examples of colocalization. Scale bars: 20 µm. (D) Quantification of colocalization between KIF3C and the indicated cargoes. Cell sample sizes: APC: 17; FMRP: 10; NgCAM: 10.

## DISCUSSION

This study addresses the function of heterodimeric Kinesin-2 family members KIF3A, KIF3B, and KIF3C in mammalian neurons. We developed a microscopy-based strategy to determine protein–protein interactions in intact neurons. With this assay, we found that KIF3AB and KIF3AC both bound KAP and that KAP recognizes the C-terminal disordered domain of KIF3s. We also refined a strategy to visualize organelle-bound KIF3AB, KIF3AC, and KAP and used that approach to generate live-cell recordings to measure the transport parameters of these organelles and to identify neuronal cargoes. KIF3/KAP organelles were mostly stationary but underwent sporadic transport, and all bind the same organelle population. Part of this population are APC-positive organelles. However, APC only accounted for ∼12 % of KIF3 organelles, indicating the existence of other cargoes. This systematic study defines the parameters of KIF3/KAP organelles in neurons and provides the tools for future work that will identify additional KIF3/KAP cargoes and determine the regulatory mechanisms of organelle-bound KIF3/KAP.

### The KIF3/KAP complex

We found that the KIF3AB/KAP complex assembles in neurons and binds organelles outside of IFT. Previous studies proposed a non-IFT function for KIF3AB in neurons, but to date there has been no evidence showing organelle-bound KIF3AB in live neurons (Takeda et al., 2000; Joseph et al., 2020; Alsabban et al., 2020; Ichinose et al., 2019, 2015; Teng et al., 2005). It is well-established that KIF3AB forms a complex with KAP in IFT. However, the structure of the assembled complex is not well defined. Early electron micrographs of KIF3AB/KAP suggested that KAP binds the C-terminal disordered domain of KIF3AB (Yamazaki et al., 1996; Wedaman et al., 1996). In contrast, biochemical analysis of interactions between heterodimeric Kinesin-2 and KAP in *Drosophila* found that KAP recognized the KIF3 coiled-coil domains (Doodhi et al., 2009; Ahmed et al., 2020). Purification of KAP for crystallography is difficult, and as a result, the structure of KIF3AB/KAP remains unsolved. We found that the C-terminal disordered domains of KIF3AB is sufficient for binding KAP, indicating that there are substantial differences in complex assembly between mammals and *Drosophila*. The clear result that the C-terminal domains of KIF3AB bind KAP may enable future studies to solve the structure of the KIF3AB/KAP and KIF3AC/KAP complexes.

A second key discovery of this study is that KIF3AC binds KAP. This interaction is mediated by the C-terminal disordered domain, analogous to the interaction between KIF3AB and KAP. Recent experiments finding that KIF3AC cannot replace KIF3AB in the IFT KIF3/KAP complex (Engelke et al., 2019) suggested that KIF3AC/KAP may not assemble, despite some biochemical evidence to the contrary (Yang and Goldstein, 1998). By showing that KIF3AC binds KAP and that KIF3C and KAP colocalize on the same organelles, our experiments resolve this seeming contradiction. Therefore, the difference between KIF3/KAP-mediated IFT and organelle transport is not due to differences in the ability of KIF3B and KIF3C to bind KAP.

### The neuronal function of KIF3/KAP

The systematic visualization of organelle-bound KIF3B, KIF3C, and KAP allowed us to quantify the trafficking behavior of KIF3/KAP cargoes and compare their “transport signature” to other kinesins. Compared to other transport vesicles, KIF3/KAP organelles moved relatively infrequently. The moving KIF3/KAP organelles exhibited velocities of 0.3 – 0.5 µm/s which are slower than those of organelles moved by Kinesin-1 or Kinesin-3 (Fu and Holzbaur, 2013; Nabb and Bentley, 2022; Yang et al., 2019; Frank et al., 2020; Farfel-Becker et al., 2019; De Pace et al., 2020), but comparable to velocities of single KIF3AB and KIF3AC motors in vitro (Guzik-Lendrum et al., 2015; Brunnbauer et al., 2010; Baumann et al., 2020; Andreasson et al., 2015; Pan et al., 2010) and during IFT in cilia (Broekhuis et al., 2014; Engelke et al., 2019). These parameters indicate that KIF3/KAP motors fulfill a specialized neuronal function that differs from other transport kinesins.

The observed trafficking behaviors are consistent with at least two functions that are not mutually exclusive. First, KIF3/KAP could act as a tether to stably maintain organelle position within neurons. The hypothesis that kinesins act as tethers is not a novel concept (Sheetz, 1987), but few examples have been described to date (Barlan and Gelfand, 2017), likely because most kinesin-related studies (including many of our own) focus on long-range transport. Microtubules in neuronal dendrites are structurally well suited for kinesin to act as tethers. Dendritic microtubules have an antiparallel organization (Baas et al., 1988; Tas et al., 2017; Weiner et al., 2021). This orientation may enable microtubule plus-end directed KIF3/KAP engaged with antiparallel microtubules to create counter-acting forces that maintain organelle positioning. A second mechanism may be that inactive KIF3/KAP binds organelles until an external signal triggers activation. The mechanisms of on-vesicle kinesin regulation are an emerging frontier in kinesin biology (Nabb and Bentley, 2022; Kumari and Ray, 2022; Verhey and Hammond, 2009; Maday et al., 2014; Cason and Holzbaur, 2022). Kinesin autoinhibition was first described for Kinesin-1 (Verhey et al., 1998) and is thought to be a regulatory mechanism for most—if not all— transport kinesins (Verhey and Hammond, 2009; Verhey et al., 2011; Twelvetrees, 2020; Chiba et al., 2022; Keren-Kaplan and Bonifacino, 2021; Bianchi et al., 2016; Ren et al., 2018; Fu and Holzbaur, 2013; Twelvetrees et al., 2019), including heterodimeric Kinesin-2 (Hammond et al., 2010). The mechanism by which organelle-bound KIF3/KAP is activated in neurons will be subject of future studies.

A key finding of the study was that KIF3AB and KIF3AC both bind KAP and localize to the same neuronal organelles. This was somewhat surprising, as KIF3AB and KIF3AC are not redundant in IFT, where KIF3C cannot compensate for a loss of KIF3B (Engelke et al., 2019). In contrast to IFT, little is known about the neuronal cargoes of KIF3/KAP. Previous studies using in vitro reconstitution (Baumann et al., 2020) and immunofluorescence (Alsabban et al., 2020) proposed APC as a KIF3AB/KAP cargo. We found robust colocalization of KIF3/KAP and APC. However, fewer than 15 % of KIF3C organelles colocalized with APC, suggesting that most KIF3/KAP organelles have a different molecular identity. Interference with KIF3/KAP causes disfunction or mislocalization of structural plasma membrane proteins (Teng et al., 2005; Soda et al., 2022; Ichinose et al., 2015) and *N*-methyl-_D_-aspartate receptors (Alsabban et al., 2020), although the causal link of these interactions is not yet fully established. Finally, KIF3/KAP organelles may function as signaling hubs that regulate neuronal development (Huangfu et al., 2003; Ichinose et al., 2019; Nishimura et al., 2004). The tools developed in this study will help elucidate the existing complexity of neuronal KIF3/KAP organelles.

### A novel assay to determine protein–protein interactions in intact neurons

To determine KIF3/KAP interactions in neurons, we developed a microscopy-based protein–protein binding assay. This approach ensures that cell-type specific accessory proteins are present. This is particularly relevant for neurons, a highly specialized cell type (Bentley and Banker, 2016; Yogev and Shen, 2017; Arendt et al., 2016, 2019). Traditional co-immunoprecipitation experiments often use non-neuronal cell types to generate sufficient protein material for biochemistry. Such experiments also rely on careful buffer calibration, as salt level and pH drastically impact protein binding. Conducting the experiment in intact neurons circumvents these problems and enables the analysis of binding interactions in the proteins’ native milieu, the cytoplasm. It also allows for multiple, relatively easy experimental replicates, as each cell functionally acts as an independent experiment.

As with any assay, proper application requires caution. Most crucially for primary neurons, interpretable experiments require cultures of consistent and high quality. This is particularly important for any colocalization experiments, as sporadic colocalization and aggresome formation tend to increase in dying cells. Targeting the candidate protein to peroxisomes with the FRB-FKBP heterodimerization system addresses this concern because it ensures that colocalization is specific and not an artifact of unhealthy cell cultures. Combined with quantitative analysis, this assay is a convenient tool for confidently defining protein–protein interactions in neurons.

## MATERIALS AND METHODS

### Cell Culture

Primary hippocampal neuronal cultures were prepared as previously described (Kaech and Banker, 2006; Kaech et al., 2012; Frank et al., 2020; Nabb and Bentley, 2022). In brief, hippocampi from E18 rats were dissected, trypsinized, dissociated, and plated on 18 mm poly-L-lysine-treated glass coverslips. Minimum essential medium with N2 supplements was used to grow cultures in a 37°C incubator with 5 % CO_2_. 6-10 DIV neurons were transfected with expression constructs and Lipofectamine 2000 (Thermo Fisher). For imaging KIF3/KAP organelles, constructs were expressed for about 3-7 h to minimize the cytosolic pool. For coexpression with candidate cargo proteins, constructs were expressed for 8-24 h to facilitate robust cargo labeling. APC required long (12+ h) expression to be visualized, likely due to its large size (2,843 residues, 310 kDa). For protein–protein binding experiments requiring expression of multiple constructs, cells were fixed after overnight expression.

### DNA Constructs

Construct details are described in Table 1. Kinesin tail construct design was previously described (Yang et al., 2019). In short, kinesin tails were designed to exclude the N-terminus containing the motor domain up to and including the first predicted coiled-coil dimerization domain (defined by Embnet).

**Methods Table 1.** Expression constructs

| Construct | Construct Design | Accession Number | Source |
| --- | --- | --- | --- |
| FRB-KIF3A tail | FRB-3myc-GGGSGGGSGGGLYK-KIF3A<sup>419-702</sup> | NM_001300792.1<br>variant 2 | Yang et al., 2019 |
| GFP-KIF3A tail | GFP-GGGSGGGSGGGLYK-KIF3A<sup>419-702</sup> | NM_001300792.1<br>variant 2 | This paper |
| Halo-KIF3A tail | Halotag-LYGGSGGGSGGGLYK-KIF3A<sup>419-702</sup> | NM_001300792.1<br>variant 2 | This paper |
| GFP-KIF3A<sup>CC</sup> | GFP-GGGSGGGSGGGLYK-KIF3A<sup>419-589</sup> | NM_001300792.1<br>variant 2 | This paper |
| Halo-KIF3A<sup>CC</sup> | Halotag-LYGGSGGGSGGGLYK-KIF3A<sup>419-589</sup> | NM_001300792.1<br>variant 2 | This paper |
| GFP-KIF3A<sup>DD</sup> | GFP-GGGSGGGSGGGLYI-KIF3A<sup>590-702</sup> | NM_001300792.1<br>variant 2 | This paper |
| Halo-KIF3A<sup>DD</sup> | Halotag-LYGGSGGGSGGGLYI-KIF3A<sup>590-702</sup> | NM_001300792.1<br>variant 2 | This paper |
| GFP-KIF3B tail | GFP-GGGSGGGSGGGLYK-KIF3B<sup>391-747</sup> | NM_008444.4 | Yang et al., 2019 |
| Halo-KIF3B tail | Halotag-LYGGSGGGSGGGLYK-KIF3B<sup>391-747</sup> | NM_008444.4 | This paper |
| GFP-KIF3B<sup>CC</sup> | GFP-GGGSGGGSGGGLYK-KIF3B<sup>391-592</sup> | NM_008444.4 | This paper |
| Halo-KIF3B<sup>CC</sup> | Halotag-LYGGSGGGSGGGLYK-KIF3B<sup>391-592</sup> | NM_008444.4 | This paper |
| GFP-KIF3B<sup>DD</sup> | GFP-GGGSGGGSGGGLYK-KIF3B<sup>593-747</sup> | NM_008444.4 | This paper |
| Halo-KIF3B<sup>DD</sup> | Halotag-LYGGSGGGSGGGLYK-KIF3B<sup>593-747</sup> | NM_008444.4 | This paper |
| GFP-KIF3C tail | GFP-GGGSGGGSGGGLYR-KIF3C<sup>411-796</sup> | NM_008445.2 | Yang et al., 2019 |
| Halo-KIF3C tail | Halotag-LYGGSGGGSGGGLYR-KIF3C<sup>411-747</sup> | NM_008445.2 | This paper |
| GFP-KIF3C<sup>CC</sup> | GFP-GGGSGGGSGGGLYR-KIF3C<sup>411-615</sup> | NM_008445.2 | This paper |
| Halo-KIF3C<sup>CC</sup> | Halotag-LYGGSGGGSGGGLYR-KIF3C<sup>411-615</sup> | NM_008445.2 | This paper |
| GFP-KIF3C<sup>DD</sup> | GFP-GGGSGGGSGGGLYT-KIF3C<sup>616-796</sup> | NM_008445.2 | This paper |
| Halo-KIF3C<sup>DD</sup> | Halotag-LYGGSGGGSGGGLYT-KIF3C<sup>616-796</sup> | NM_008445.2 | This paper |
| FRB-KAP | FRB-3myc-GGGSGGGSGGGLYK-KAP3B | NM_001375831.1 | This paper |
| Halo-KAP | Halotag-LYGGSGGGSGGGLYK-KAP3B | NM_001375831.1 | This paper |
| GFP-KIF5C tail | GFP-GGGSGGGSGGGLYKGGGSGGGSGGGP-KIF5C<sup>378-955</sup> | NM_001107730 | Yang et al., 2019 |
| PEX3-tdTomato-FKBP | PEX3-GGSGSGGPRPT-tdTomato-RTGASGGGSGGGSGGGSGGGTR-FKBP | NM_003630.3 | This paper |
| APC-mEmerald | APC-FGGASAASGSADPPVAT-mEmerald | NM_000038.6 | Adapted from Morin et al., 1997 |
| GFP-FMRP | GFP-SGLRSRAQASNSMEDYKDDDDK-FMRP | NM_002024.6 | Adapted from Stetler et al., 2006 |
| NgCAM-GFP | NgCAM-KLGAPRPT-GFP | NM_205153.1 | Nabb and Bentley, 2022 |

### Imaging

Live-cell recordings were acquired with an Andor Dragonfly built on a Nikon Ti2 with a CFI Apo 60x 1.49 objective and two sCMOS cameras (Zyla 4.2+; Andor). The imaging stage, microscope objective, and sample were maintained at 37°C in a full lexan incubation ensemble (OkoLab). The z-axis movement was controlled by a Perfect Focus System. During imaging, neurons were maintained in Hibernate E without phenol red (BrainBits). Images were acquired at 2 frames per second for 30 s. Axons were identified with an anti-neurofascin antibody (NeuroMab Cat# 75-027) conjugated to CF405 (Mix-n-Stain CF405S antibody labeling kit; Biotum Cat# 92231) in the imaging medium.

For fixed-cell recordings, neurons were fixed with 4 % paraformaldehyde/4 % sucrose in phosphate-buffered saline for 20 min at 37°C. Coverslips were mounted on slides with ProLong Diamond Anti-fade (Thermo Fisher Cat# P36961). Fixed cell images were recorded with a Zeiss Axio Imager Z1 microscope equipped with a 63x 1.4 Plan-Apochromat objective and a CCD (Axiocam 506 mono; Zeiss).

Halotag-expressing cells were incubated with 50 nM JF549 or JF646 (Grimm et al., 2015) for 5 min and washed with conditioned medium for 5 min prior to imaging or fixation. For protein– protein binding assays, neurons were incubated with 100 nM linker (AP21967; Takara, Cat# 635056) for 2 h before fixation.

### Analysis

All analysis was performed by a single blinded reviewer. To ensure consistency and reduce bias, multiple experiments were combined into a large dataset and the analyst was blinded to the condition.

Kymographs were generated with MetaMorph software (Molecular Devices). A single analyst manually identified all transport events from kymographs. Each continuous line with a constant slope was scored as a single transport event. A single organelle could undergo multiple transport events if there was a distinct pause (>3 frames) between them. Event coordinates were exported to Microsoft Excel or GraphPad Prism for statistical analysis. To ensure that only microtubule-based long-range transport events were included in the analysis, excursions <2 µm were excluded. Run length and velocities were calculated with 1 µm bin size and a 0.1 µm/s bin sizes, respectively. Mean run lengths were calculated using a single exponential decay following the equation

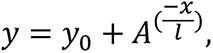

where *A* is the maximum amplitude, and *l* is the mean run length reported as ±SEM. The velocity was curve fit to a log-normal distribution, following the equation

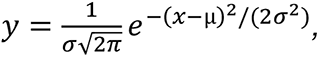

where µ ∈ (−∞, +∞) and σ > 0, and the geometric mean following the equation

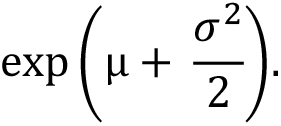

The percentage of stationary organelles and the total number of organelles were generated by manual counting in the same regions of axons and dendrites used for kymograph analysis. Percent stationary organelles was calculated from the number of moving organelles (detected by kymograph analysis) the number of total organelles in these same regions.

To calculate the Pearson’s correlation coefficients for protein–protein binding experiments (Fig. 1-4), three to five clustered and brightly-labeled peroxisomes were identified in a dendrite. A region of interest (ROI) was drawn around these peroxisomes, and the Pearson’s correlation coefficient was calculated with the Coloc2 FIJI plugin.

For KIF3 organelle colocalization (Fig. 7) Pearson’s correlation coefficients were generated from representative dendrites of each neuron.

The percent colocalization of KIF3C with cargoes (Fig. 8) was calculated by manual counting of fluorescent structures in both channels. Organelles were classified as “not colocalized” and “colocalized with comparable morphology”.

P values for Pearson’s correlation coefficients were determined by Student’s t-test. Error bars show the standard error of the mean.

## Abbreviations used

KAP: kinesin-associated protein 3B
IFT: intraflagellar transport
APC: adenomatous polyposis coli
FMRP: fragile-X-mental retardation protein
NgCAM: neuron-glia cell adhesion molecule

## ACKNOWLEDGMENTS

We thank Geraldine Quinones for excellent technical assistance in culturing and maintaining hippocampal neurons. We thank Jasper Jeffrey for help with figure design. We thank the BioResearch facility at Rensselaer for assistance with husbandry and tissue collection. Research reported in this publication was supported by the National Institute of Mental Health award R01-MH066179 to M.B., National Institute of General Medical Sciences award R37-GM054141 to S.P.G, and National Institute of General Medical Sciences Training Program T32GM067545 predoctoral fellowship to A.G. The content is solely the responsibility of the authors and does not necessarily represent the official views of the National Institutes of Health.

## AUTHOR CONTRIBUTIONS

A.G. and A.M. designed and performed experiments, analyzed data, and wrote the manuscript. S.P.G. and M.B. supervised the research, conceptualized the study, designed experiments, and wrote the manuscript.

